# Transcriptome-Driven Constraint-Based Modelling Reveals Metabolic Targets for Ovarian Cancer

**DOI:** 10.1101/2025.06.24.661329

**Authors:** Kate E. Meeson, Joanne McGrail, Jean-Marc Schwartz, Stephen S. Taylor

## Abstract

Constraint-based modelling (CBM) is a powerful computational approach that reconstructs cellular metabolism by integrating ‘omics data with genome-scale metabolic models (GEMs), enabling *in silico* hypothesis generation and genetic engineering studies. Advances in high-throughput ‘omics technologies and the complete mapping of the human genome have expanded the application of CBM to human systems. Given that altered metabolism is a hallmark of cancer, this disease represents an ideal context for developing and applying CBM workflows. Despite the presence of well-characterised metabolic signatures and vulnerabilities in ovarian cancer, this tumour type remains under-explored within the CBM field. Meanwhile, the limited efficacy of current therapies and the frequent emergence of chemoresistance underscore the need for novel, mechanism-based approaches to therapeutic discovery. In this study, we constructed ovarian cancer-specific metabolic models using an ‘omics integration algorithm that incorporates transcriptomic data in a way that is directed by experimental growth measurements. Simulations identified multiple candidate molecules predicted to influence cancer cell proliferation. Among these, triosephosphate isomerase 1 (TPI1) was selected for experimental validation based on qualitative prioritisation criteria. Notably, model predictions were supported by RNA sequencing and proliferation assays, implicating TPI1 in ovarian cancer cell growth. Our results provide novel insights into the metabolic dependencies of ovarian cancer and demonstrate a multi-omics CBM workflow that may be broadly applicable for uncovering therapeutic vulnerabilities in other malignancies.

## INTRODUCTION

Ovarian cancer ranks as one the most common and fatal gynaecological cancers. Around 4.4% of all cancer-related deaths in women are attributed to ovarian cancer (Momenimovahed et al., 2019) and as a result of late diagnosis, the 5-year survival rate is relatively low at 29% (Reid et al., 2017). The majority of tumours (90%) are epithelial in origin and are organised into five distinct histological subtypes, all with independent clinical phenotypes and developmental mechanisms (Momenimovahed et al., 2019; Reid et al., 2017). As a first-line treatment, patients undergo cytoreductive surgical debulking, alongside platinum-based chemotherapy (Cortez et al., 2018). Other treatment avenues include, but are not limited, to angiogenesis inhibitors (e.g. bevacizumab) and PARP1/2 inhibitors for patients with homologous recombination deficient tumours (Cortez et al., 2018). Despite various treatments being available for ovarian cancer, their efficacy is hindered by chemoresistance, which results in at least a 60% risk of recurrence (Cortez et al., 2018).

Within the past couple of decades, our understanding of the metabolic reprogramming of cancer cells has improved, allowing us to formulate explanations for clinical phenotypes. Cellular metabolism can be organised into distinct metabolic subsystems and ovarian cancer shows dysregulation across many of these, including drug metabolism, energy metabolism, glucose metabolism, amino acid metabolism and lipid metabolism. For example, the activity of ABC drug transporters, of which *ABCB1* is overexpressed in ∼20% of relapsed high-grade serous ovarian cancers (Christie et al., 2019), is underpinned by mitochondrial respiration (Giddings et al., 2021; Robey et al., 2018). A fundamental aspect of cancer metabolism is the rewiring of energy metabolism, which is now a well-recognised hallmark of cancer, supporting one of the original cancer hallmarks: sustained proliferative potential (Hanahan & Weinberg, 2011; Pavlova & Thompson, 2016). This is especially true for ovarian cancer, where metabolic signatures are being proposed. For example, increased rates of oxidative phosphorylation have been correlated with a higher invasive potential (Nantasupha et al., 2021). Furthermore, underlying signalling pathways have been linked to these observations, for example mutations across the PI3K/Akt pathway have been shown to disrupt the expression of the GLUT1 transporter and glycolytic enzymes (Pavlova & Thompson, 2016). In addition, amino acid signatures associated with ovarian cancer have been proposed, including elevated tryptophan metabolism and higher levels of phenylalanine and tyrosine-derived metabolites in patient ascites (Grobben et al., 2023). Furthermore, ovarian tumours hijack lipid metabolism to fuel cellular proliferation (Chaudhry et al., 2022; Ji et al., 2020). Given this knowledge, it is important to improve our understanding of ovarian cancer metabolism to reveal mechanisms driving clinical treatment response and to highlight potential enzymatic targets.

One approach to studying the metabolism of cancer cells is constraint-based modelling, which involves computationally reconstructing the metabolic network of a cell to generate a genome-scale model (GEM). GEMs can be personalised for a specific cell type or disease through the integration of enzyme expression data generated from that same cell type or disease. For each enzyme-regulated reaction in a GEM, there is a gene-protein-reaction (GPR) rule that dictates how the enzyme expression data is integrated. Through the integration of enzyme expression data, for example gene or protein expression values, the user can place boundaries on the reaction rates (fluxes), which in turn affects the predicted flux values for downstream metabolic reactions. The integration of ‘omics data in this way transforms the unconstrained, solution space to a more realistic, biologically relevant representation, from which a flux distribution can be estimated using flux balance analysis (FBA) (Orth et al., 2010). These sample-specific constraint-based models serve as a framework for *in silico* genetic engineering of targets to predict the ‘real-world’ outcome, where the absence or overexpression of a particular enzyme can be simulated.

Relative to other cancers, the constraint-based modelling of ovarian cancer is limited, however, a few studies provide direction and support for further investigation. The earliest example is an investigation into cisplatin resistance in the A2780 ovarian cancer cell line, where transcriptomics was integrated with the Recon1 human GEM (Motamedian et al., 2015). Another study used constraint-based modelling to explore a metabolic basis for the metastatic transition of the OVCAR3 cell line, with subsequent *in vitro* work to confirm drug target predictions (Arora et al., 2023). Finally, in the development of the algorithm which will be used to integrate transcriptomics in this study, the metabolic flux distributions of low- and high-grade serous ovarian cancer were predicted and compared, with validation against publicly available CRISPR-Cas9 data (Meeson & Schwartz, 2024).

Here, the recently developed single-omics integration algorithm (Meeson & Schwartz, 2024) has been used to constrain numerous ovarian cancer-specific metabolic models using gene expression data. These models were then used as scaffolds for gene deletion simulations to identify a suitable target for *in vitro* knockdown to test the model-predicted inhibition of cell line growth. Growth studies and RNA sequencing (RNAseq) analysis were used to validate modelling predictions and disentangle signalling pathways underpinning ovarian cancer cell dependency on TPI1, in turn presenting a workflow that has been experimentally validated and could be translated to other cancer studies.

## RESULTS

### Using constraint-based models as a scaffold for gene knockdown predictions

To enable computational simulations and facilitate the understanding of specific ovarian cancer cell line metabolism, we constrained 46 individual metabolic models using publicly available gene expression data. Transcriptomics from the Cancer Cell Line Encyclopedia (CCLE) (Barretina et al., 2012; Cerami et al., 2012; Gao et al., 2013), which were specific to individual cell lines, were integrated with the gene-protein-reaction rules in the Human1 GEM using a recently designed algorithm (Meeson & Schwartz, 2024). To verify that these models were predicting experimentally realistic phenotypes, we then compared the FBA-predicted growth rates with experimental doubling times from the CCLE (Supplementary file ‘CCLE_2020_transcriptomics_annotations.xlsx’) (***Figure 1****a*). This comparison informed a significantly positive correlation (r) between experimental growth rates and constraint-based modelling predictions (0.5386) (p<0.0001), confirming that these metabolic models were able to recapitulate the growth rate of in vitro cell lines. This agreement between models and experimental measurements provided incentive to explore these models as gene engineering platforms to study growth inhibition of ovarian cancer cell lines.

**Figure 1.**
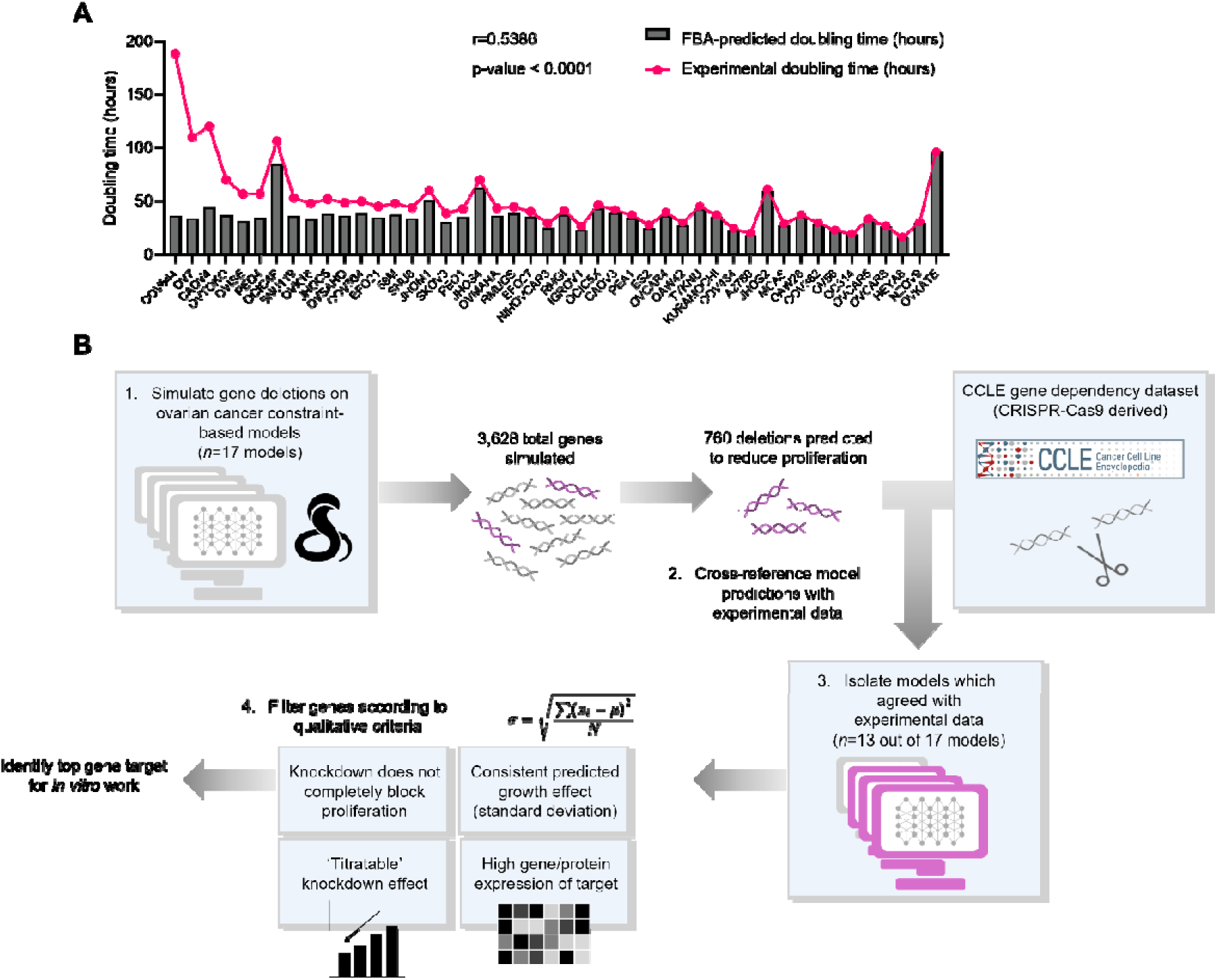
Constraint-based models can be used as accurate predictive platforms. A) Comparison between experimental doubling times (pink) and constraint-based modelling predicted doubling times (grey) for 46 ovarian cancer cell line-specific models. CCLE transcriptomics has been used to constrain models. Bars organized with the greatest difference between experimental and model-predicted on the left-hand side, and the smallest difference on the right-hand side. Pearson correlation coefficient (r) between experimental and predicted was 0.5386 (p-value<0.0001). B) Graphic for the workflow for the prediction of gene knockdown targets using constraint-based models.

Once the metabolic models had been constrained and had their growth predictions validated, we needed to determine how accurately they could predict the result of gene engineering experiments. Therefore, we used the models to predict the essentiality of a panel of genes for cell proliferation, by simulating the deletion of these genes across the models. This gene deletion was mimicked by simulating an expression value of zero in place of a positive gene expression measurement, or in place of a previously unconstrained enzyme-regulated reaction. Alongside the transcriptomics that we used to formulate model constraints, the CCLE have made a CRISPR-Cas9 gene dependency (DepMap) dataset publicly available (https://doi.org/10.25452/figshare.plus.24667905.v2; ‘CRISPRGeneDependency.csv’) (Tsherniak et al., 2017), which describes the essentiality of thousands of genes across the human genome, based on cell line proliferation before and after a gene knockout. This experimental dataset, in combination with our constraint-based models, allowed us to develop a pipeline to identify a single gene target for knockdown to reduce the proliferation of an ovarian cancer cell line (***Figure 1b***).

There are 3,628 total enzyme-encoding genes across the Human1 metabolic model, therefore the deletion of these genes was simulated and compared with the CCLE experimental dataset. Of these 3,628 total genes, simulations indicated that the deletion of 760 genes would reduce cell proliferation (defined as a reduction in predicted growth rate of at least 5% compared with the original cell line-specific model prediction). There were 715 out of these 760 model genes that were also present in the CCLE dataset, therefore the *in silico* ‘growth effect’ (growth rate predicted after deletion/original predicted growth rate) was correlated to the corresponding experimental value for these 715 genes. This analysis showed that there was a significant correlation between model predictions and CCLE experimental gene dependency scores for 13 out of 17 models evaluated, indicating that these 13 models could accurately predict the growth effect of a gene deletion. The range of correlation coefficients (r) achieved by these 13 models ranged from -0.118 to -0.500 (p˂0.05), with a mean of -0.288 (Supplementary Table 2).

To filter this subset of 715 genes further and identify an individual target for experimental exploration, four criteria were applied (***Figure 1b***): 1) a consistent growth effect across the 13 models (as measured by standard deviation); 2) gene knockdown would not completely inhibit cell growth as this would hinder sample preparation for RNA sequencing analysis; 3) gene effect is ‘titratable’, and therefore predicted to cause inhibition upon partial knockdown as well as full gene knockout; and 4) the protein encoded by this gene is relatively highly expressed across ovarian cell lines.

To further narrow down this list of 715 candidate genes, more detailed modelling simulations were run to predict the effect of an siRNA-mediated knockdown. There were 13 out of 17 models that significantly correlated with CCLE experimental growth dependency scores (p˂0.05), and only these 13 models were allowed to proceed to further simulations.

Here, we aimed to identify a gene target that would represent a potential metabolic vulnerability across as many different ovarian cancer cell lines as possible. Therefore, we studied consistency in predicted knockdown effect across the 13 cell line-specific models. Our measure of this consistency was the standard deviation in predicted growth effect (growth rate predicted after deletion/original predicted growth rate) across these 13 models (***Figure 1b***). At a 30% standard deviation in predicted growth effect, there were 53 potential gene targets (out of the initial 715 subset). When this threshold was lowered to 27% standard deviation, there were eight potential targets for *in vitro* growth inhibition.

Next, we wanted to understand whether the growth dependency of the cell line-specific models on these eight targets was ‘titratable’. Importantly, a ‘titratable’ growth dependency would mean a full *in vitro* gene deletion need not be achieved as models would predict growth inhibition at a partial knockdown – as is accomplished through siRNA-mediated transfection. To select the initial target list, a full deletion had been simulated, by setting both the lower and upper bounds of the enzyme-regulated reaction fluxes to 0 mmol/gDW/hour across the models. To simulate a partial gene knockdown, the reaction bounds of these enzyme-regulated reactions were gradually relaxed to allow more metabolic flux to flow through. Following this partial knockdown simulation, we concluded that of the eight potential targets, four remained: triose phosphate isomerase 1 gene (*TPI1*) and three genes encoding the ATP synthase membrane complex (*ATP5MC2*, *ATP5MC3* and *ATP5MK*) (***Figure 2a***).

**Figure 2.**
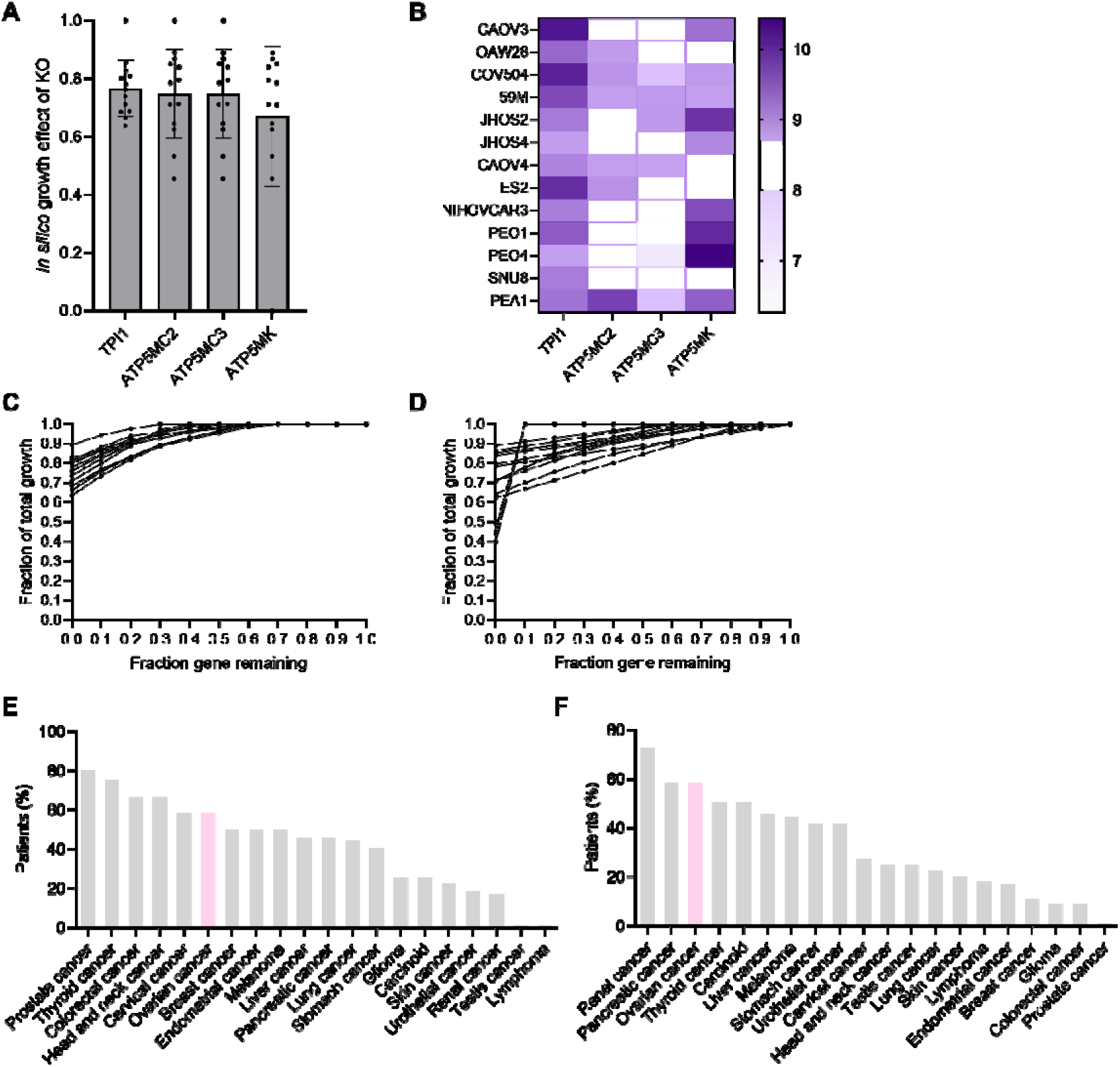
Modelling predicts triosephosphate isomerase 1 as a target for *in vitro* growth inhibition. A) Growth effect of the top four knockout targets from an *in silico* knockout simulation (ratio of growth rate after/before knockout). Individual data points correspond to individual models, and the bars show standard deviation. Simulations ran on the 13 models that significantly correlated with CCLE gene dependency dataset. B) Heatmap for the gene expression of four target genes across 13 ovarian cancer cell lines. CCLE 2020 transcriptomics. C) Model simulation of the growth effect of TPI1 knockdown, where the bounds of the TPI1-catalysed Human1 reaction (MAR04391) have been constrained to 0–100% of the original bounds. D) Model simulation of growth effect of ATP5MK/C2/C3 knockdown on n=13 models, where the lower and upper bounds of the ATP synthase-catalysed Human1 reaction (MAR06916) have been constrained. E) ATP5MC2 protein expression results, from Human Protein Atlas. Percentage of patient tumour samples with high or medium protein expression, categorised for pathology. Antibody dataset HPA051469, accessed 11/10/2024. F) TPI1 protein expression results, from Human Protein Atlas. Percentage of patient tumour samples with high or medium protein expression, categorised for pathology. Antibody dataset HPA053568, accessed 11/10/2024.

The same selection criteria as described above were used to evaluate TPI1 and the ATP synthase membrane complex-encoding genes. When a full gene deletion was simulated across cell line-specific models, TPI1 showed better consistency than the ATP synthase targets, with a predicted growth effect standard deviation of less than 10%, compared with 15% and 24% for *ATP5MC2/3* and *ATP5MK*, respectively **(*Figure 2a***). In addition, in the CCLE dataset, which was used to constrain models, *TPI1* has higher expression than the ATP synthase targets (***Figure 2b***). Furthermore, models predicted that *TPI1* had the most ‘titratable’ knockdown phenotype, as indicated by a smoother decline in growth rate as a greater proportion of the gene was knocked down *in silico* (***Figure 2c*** compared with ***Figure 2d***). Finally, protein expression data from the Human Protein Atlas (HPA) (*Human Protein Atlas*, 2024; Uhlén et al., 2015) suggested that there is relatively high expression of the TPI1 protein in ovarian cancer patient-derived tumour samples, relative to other tissue types. Namely, of 20 different cancers, ovarian cancer has the third highest proportion of tumours with medium to high levels of TPI1 expression, compared with the fifth highest proportion for ATP5MC2 (*Figure 2e* and *Figure 2f)*.

In conclusion, cell line-specific models suggested *TPI1* as the most promising candidate for experimental growth inhibition studies according to a range of criteria, including a consistent growth effect across cell lines, high gene and protein expression and a titratable knockdown phenotype. Therefore, *TPI1* proceeded to *in vitro* studies to simultaneously validate model predictions and explore the role of the TPI1 enzyme across ovarian cancer metabolism.

### TPI1 impacts the colony formation and proliferation of an ovarian cancer cell line

Once the constraint-based models had been shown to accurately predict experimental phenotypes, namely growth rate and the growth dependency of a panel of metabolic genes, we wanted to explore the role of TPI1 in an ovarian cancer cell line, since models predicted TPI1 to be a worthwhile target for growth inhibition. To explore the function of this protein in ovarian cancer, we used siRNA-mediated knockdown of the *TPI1* gene and colony formation and proliferation assays.

Experimental work was performed in the OV56 cell line, firstly because there was high similarity between experimental and model-predicted growth rates (***Figure 1***a) and secondly, because OV56 was not included in the target prediction (***Figure 2***), therefore, this would test the broader applicability of model predictions. There is limited evidence for the role of TPI1 in ovarian cancer, however, it has recently been reported that *TPI1* knockdown in the A549 lung adenocarcinoma cell line reduced colony formation potential and cell proliferation (P. Liu et al., 2022), so we included the A549 cell line as a comparison.

We wanted to understand how the role of TPI1 in OV56 compared with an oncogene known to contribute to growth dysregulation in ovarian cancer. One such oncogene is *MYC*, which has been implicated in virtually all stages of tumorigenesis, including altered metabolism, via transcriptional activation (H. Chen et al., 2018; Dang, 2012). MYC is amplified at the gene and protein level across ovarian cancers, with associations to chemoresistance, the prevention of cell cycle arrest and increased activation of apoptosis, although MYC is often described as ‘undruggable’ since its oncogenic activity is through transcriptional regulation rather than direct enzymatic activity (Reyes-González & Vivas-Mejía, 2021). Therefore, we compared the effect of siRNA targeting *TPI1* (siTPI1) or *MYC* (siMYC) on the colony formation potential and short-term cellular proliferation of OV56 and A549, to position TPI1 as a potential drug target for future studies.

Prior to colony formation and cell proliferation assays, it was necessary to confirm the successful knockdown of *TPI1* in the OV56 cell line. Using siTPI1 transfection, immunoblotting confirmed a knockdown to approximately 10% of the initial protein expression levels in the OV56 cell line (***Figure 3****a*). Across a 10-day colony formation assay, *TPI1* knockdown in the OV56 cell line resulted in smaller and fewer colonies (***Figure 3****b*), which equated to approximately 20% of the survival of the non-targeting siRNA (NT siRNA) control, with statistical significance (***Figure 3****c*). In the A549 lung adenocarcinoma cell line, *TPI1* knockdown reduced colony formation potential (***Figure 3****d*), which indicated significance when quantified (***Figure 3****e*). The level to which siTPI1 reduced colony formation in the OV56 and A549 cell lines was comparable to siMYC (***Figure 3b*** and ***Figure 3d***).

**Figure 3.**
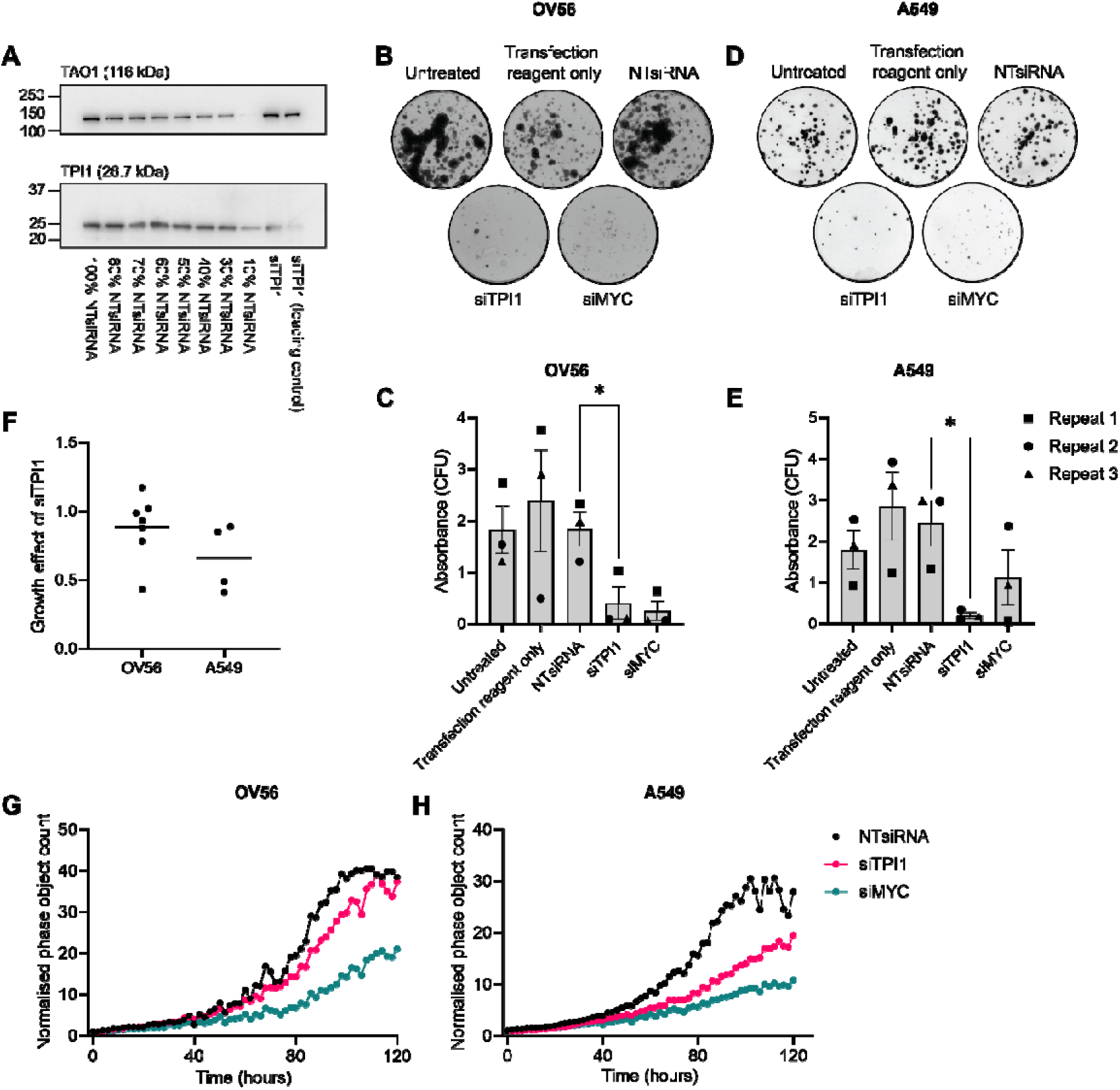
TPI1 knockdown reduces colony formation ability and cell proliferation of OV56 and A549. A) Representative immunoblot for TPI1 in the OV56 cell line. Non-targeting siRNA (NTsiRNA) has been titrated to allow an approximate estimate of the TPI1 protein expression in the siTPI1 sample. B) Exemplar colony formation assay for the OV56 cell line transfected with siRNA (NTsiRNA, siTPI1 or siMYC) or exposed to negative control conditions (untreated or transfection reagent only). C) Quantification of colony formation assays. *p<0.05. Bars indicate standard error. Values have been normalised to NTsiRNA. Three biological replicates represented by data point shapes. D) Exemplar colony formation assay for the A549 cell line transfected with siRNAs as in B. E) Quantification of the colony formation assays. *p<0.05. Bars indicate standard error. Values have been normalised to NTsiRNA. Three biological replicates represented by data point shapes. F) Growth effect of siTPI1 (OV56 and A549 cell lines) (calculated as doubling time before/after siRNA transfection) compared with NTsiRNA calculated as a ratio of the normalised phase object count at time point 100 hours for the siTPI1 compared with the NTsiRNA conditions. Biological replicates and mean growth effect have been plotted. G) Representative plot for the growth of OV56 cells upon siRNA transfection (NTsiRNA: black, siTPI1: pink, siMYC: teal). Live cell microscopy assay, using the Incucyte S3^®^ live-cell analysis system (Sartorius). Biological repeats for this experiment have been shown in Supplementary Figure 1. H) Representative plot for the growth of A549 cells upon siRNA transfection. Live cell microscopy assay, using the Incucyte S3^®^ live-cell analysis system (Sartorius). Biological repeats for this experiment have been shown in Supplementary Figure 1.

To determine the role of the TPI1 enzyme in cellular proliferation, live-cell imaging was used to study OV56 and A549 over 4-day culture, following siTPI1 or siMYC transfection. As a measure of growth inhibition, a mean growth ratio (doubling time before/after siRNA transfection) was calculated. For the OV56 cell line, this growth ratio was variable, and for some biological replicates, was above 1.0, indicating siTPI1 could increase the rate of proliferation of the OV56 cell line (***Figure 3f***). However, the resounding effect of siTPI1 transfection was a reduction in cellular proliferation across both the OV56 and A549 cell lines, with mean growth ratios of 0.89 and 0.66, respectively (***Figure 3f***). For both cell lines, a reduction in proliferation could be observed past 80 hours of culture with siTPI1 or siMYC transfection, suggesting *TPI1* regulates cell proliferation in a similar manner to the *MYC* oncogene (***Figure 3****g* and ***Figure 3****h*). In conclusion, experimental results support the dependency on expression of the TPI1 enzyme for OV56 cell line growth, which confirms predictions made using constraint-based modelling.

### RNA sequencing suggests that expression of TPI1 promotes proliferation, migration and survival of an ovarian cancer cell line

Following experimental work confirming dependency on *TPI1* for proliferation and colony formation of the OV56 cell line, an RNA sequencing (RNAseq) analysis was performed to identify the genes and pathways driving this dependency. Gene expression results confirmed the successful knockdown of *TPI1*, where only 14% of transcripts remained in the *TPI1* knockdown sample, compared with the NT siRNA control (***Figure 4***a). When visualised using PCA (***Figure 4****b*) and heat mapping (***Figure 4****c*), it could be observed that the knockdown of *TPI1* greatly disrupted the gene expression of OV56, where knockdown samples were distinct from all negative controls across three biological repeats.

**Figure 4.**
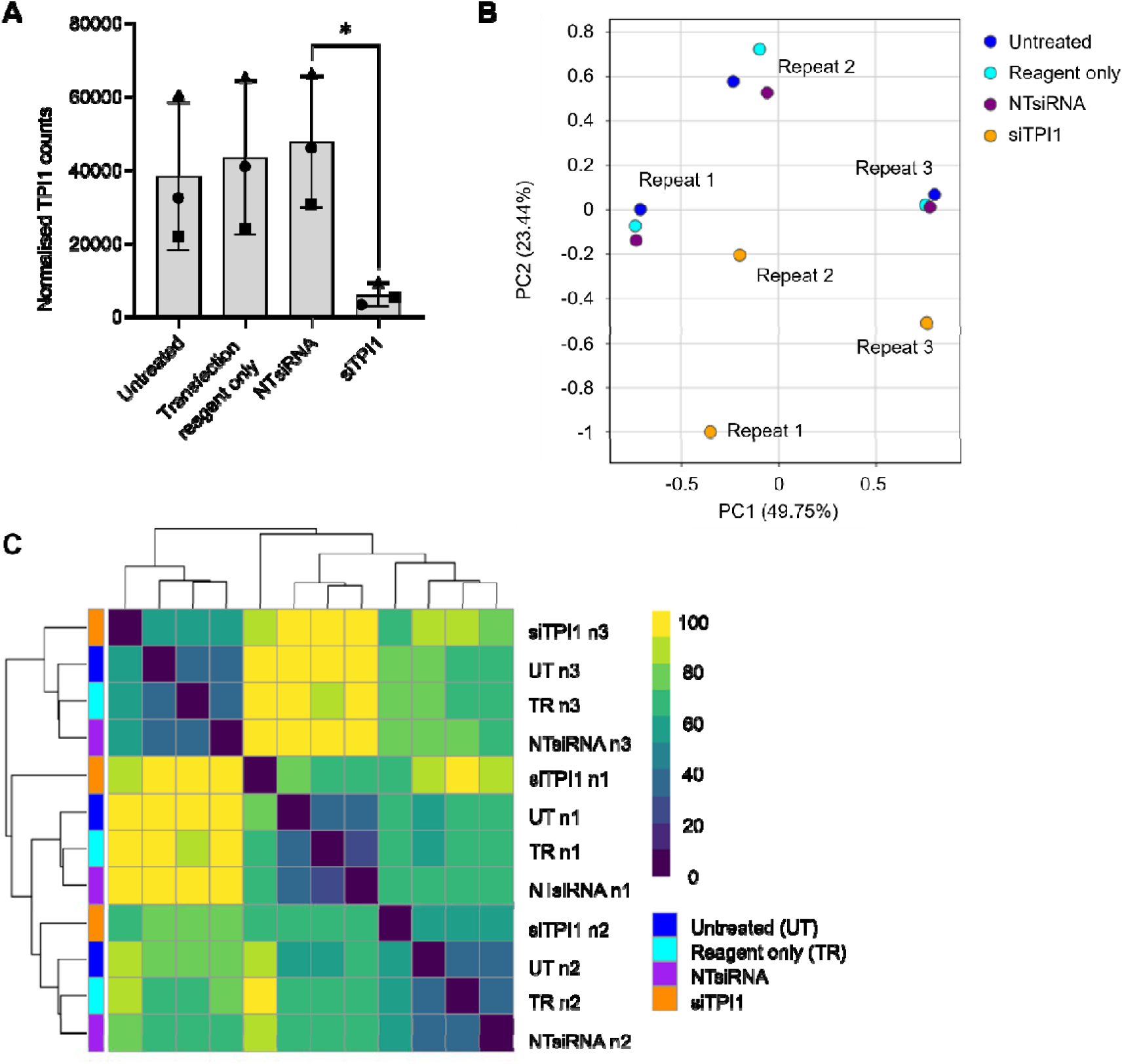
TPI1 knockdown regulates gene expression of OV56. A) Normalised gene expression of TPI1 across RNAseq sample conditions (x-axis). Biological repeats are labelled with symbols. Rlog normalised gene counts. Asterisk indicates statistical significance (p=0.0451; paired t-test). Bars indicate standard error. B) PCA of rlog-transformed normalised counts, from DESeq2. Biological repeats and siRNA conditions indicated on figure. C) Heatmap of rlog-transformed normalised counts. Biological repeats and siRNA conditions indicated on figure.

Differential gene expression analysis was performed to determine those genes showing the greatest degree of change in expression upon siTPI1 transfection. There were 2,419 genes upregulated, and 2,584 genes downregulated in the siTPI1 condition compared with NT siRNA control (FDR<0.05 and Log2FC>0) (***Figure 5****a*). In total, 15% of the initial 33,253 nonzero read count subset was differentially regulated in OV56 upon *TPI1* knockdown. According to their FDR values, the top ten differentially regulated genes included *EMC10*, *PLBD2*, *SIGMAR1*, *MX1*, *TPI1P1*, *PFN1*, *DDAH1*, *HMGA1*, *TMEM9* and *TAGLN2* (***Figure 5****a*). In comparison to the fold-change observed for *TPI1*, which was downregulated by 0.13-fold in the siTPI1 sample relative to the NT siRNA control (padj=1×10⁻⁴²), the fold-changes of the aforementioned most differentially regulated genes ranged from 0.046 to 4.5 (***Figure 5****a*). In general, the knockdown of *TPI1* caused the subsequent downregulation of genes in OV56 cells, with 44 out of the 50 top differentially expressed genes having been downregulated in the siTPI1 condition (***Figure 5****b*).

**Figure 5.**
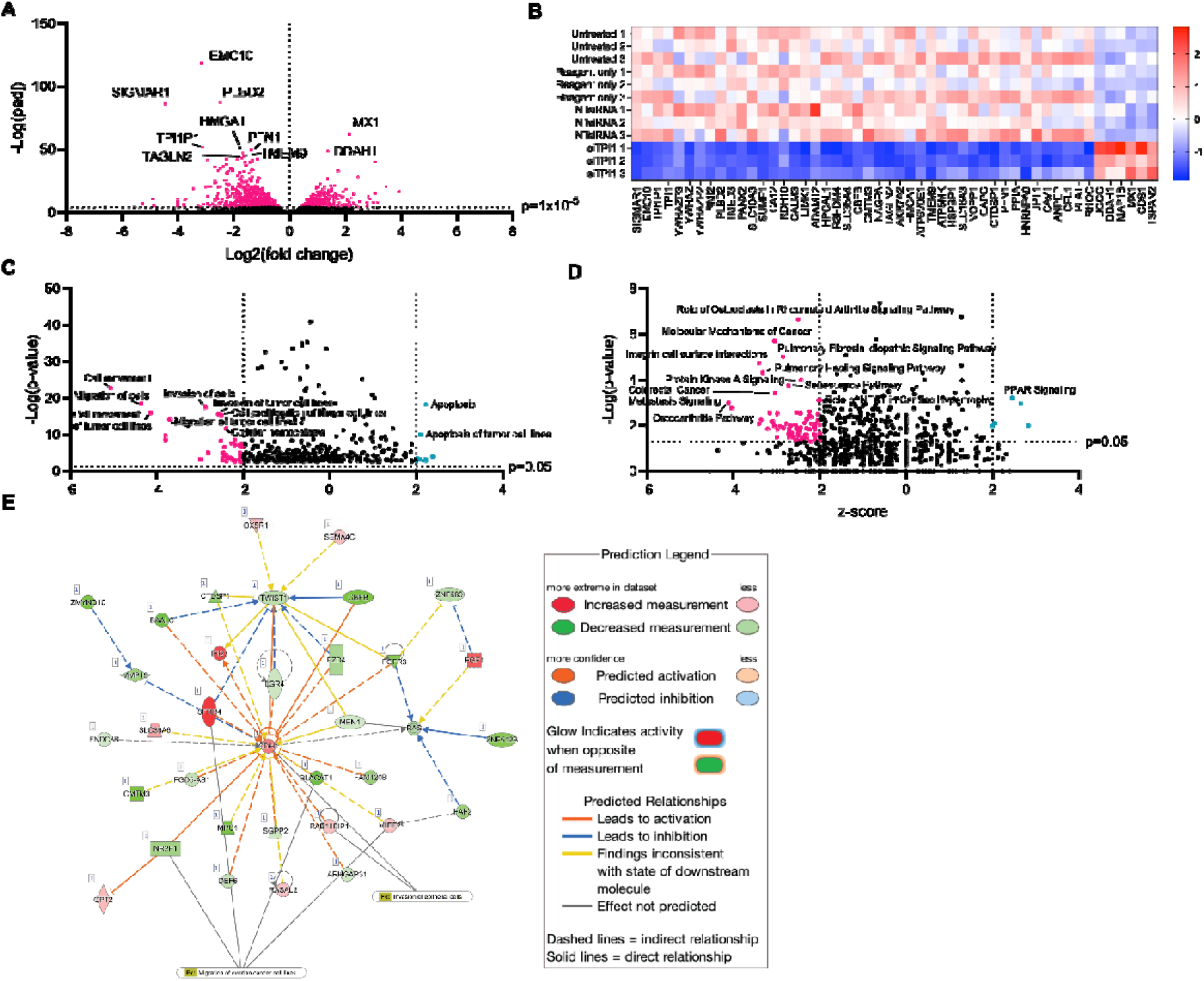
Differentially expressed genes and pathways upon TPI1 knockdown. A) Volcano plot for the RNAseq comparison between NTsiRNA and siTPI1 conditions, in OV56 cells. Log2(fold change) (Log2FC) has been plotted against -Log(adjusted p-value). The adjusted p-value threshold was 1×10^-5^ (pink data points). The top ten hits have been labelled. B) Heatmap for the top 50 differentially expressed genes between NTsiRNA and siTPI1. Top hits were selected according to p-values reported from DESeq2 analysis. Genes have been ordered according to fold-change along the x-axis. Values shown are the z-scores ((x-µ)/ σ), calculated from the rlog-normalised gene expression (upregulation indicated by red cell, downregulation indicated by blue cell). C) Volcano plot for the biological functions associated with differentially expressed genes between NTsiRNA and siTPI1 conditions. Analysis performed using QIAGEN IPA. Z-score was calculated to statistically compare input gene expression to literature predictions (thresholds of >+2 or <-2 indicate activation or inhibition). -Log(p-value) threshold was set to 1.3, as this is equal to a p=0.05. Pink: process predicted to be inhibited; teal: predicted to be activated. Top ten processes have been labelled. D) Canonical pathway annotation of differentially expressed genes. Same parameters as in C. E) Network of proliferation-associated genes with differential expression upon TPI1 knockdown. Analysis performed using QIAGEN IPA. The definition of colour of edges and nodes, as well as dashed/solid lines has been indicated on key. Network 4 from DESeq2 analysis, which involves cellular movement processes. Connections to ‘invasion of epithelial cells’ and ‘migration of ovarian cancer cell lines’ have been overlayed.

Functional enrichment showed that the knockdown of *TPI1* was associated with specific biological functions and canonical signalling pathways. Our experimental evidence suggested that cellular proliferation and colony formation of OV56 is dependent on the TPI1 enzyme (*Figure 3*), and in agreement, RNAseq analysis highlighted proliferation, migration, movement and invasion as being inhibited upon *TPI1* knockdown (***Figure 5****c*). In addition, cellular homeostasis was downregulated in the siTPI1 condition (***Figure 5****c*), indicating that the knockdown of *TPI1* disrupts the stability of the internal environment of the OV56 cell line. Concordant with published results obtained in Sertoli cells exposed to miRNA-mediated *TPI1* inhibition (An et al., 2022), functional annotation suggested that the knockdown of *TPI1* in OV56 promoted cellular apoptosis, and ‘apoptosis of tumour cell lines’ (***Figure 5****c*).

Regarding the regulation of canonical signalling pathways, RNAseq analysis suggested that the knockdown of *TPI1* could downregulate as many as 76 pathways and activate five pathways (***Figure 5****d*). These could provide a molecular mechanism for the reduced cellular proliferation upon *TPI1* knockdown, for example, integrin cell surface interactions were predicted to be downregulated (***Figure 5****d*), which could inhibit the binding of OV56 to the extracellular matrix and subsequently limit cell growth. Furthermore, pathway analysis predicted the downregulation of protein kinase A signalling upon *TPI1* knockdown (***Figure 5****d*), and this could relate to apoptosis as Protein Kinase A promotes cell survival in response to glucose starvation via metabolic rewiring (Palorini et al., 2016). RNAseq analysis also indicated that *TPI1* knockdown activates the PPAR signalling pathway (***Figure 5****d*), which is involved in glucose and lipid metabolism, and activation of PPARᵧ has been proposed as a therapeutic strategy for ovarian cancer (Vignati et al., 2006).

Gene network analysis was performed to understand the individual gene-gene interactions that drive the biological functions of TPI1 in OV56. Both colony-formation and proliferation assays and functional enrichment of gene expression data indicated that the knockdown of *TPI1* reduces cell proliferation, therefore a gene network describing the proliferative effects of TPI1 knockdown was generated (***Figure 5****e*). Network analysis reported statistically significant connections to ‘invasion of epithelial cells’ (p=0.00142) and ‘migration of ovarian cancer cells’ (p=5.31×10^-5^), via multiple genes (***Figure 5****e*). Results indicated there was a ‘hub’ of gene interactions centred on the cadherin-1 (CDH1/E-cadherin) gene, which was upregulated by 1.05-fold in the siTPI1 sample (padj=0.000927) (***Figure 5****e*).

In conclusion, RNAseq analysis predicted a molecular basis for the growth inhibition that we observed experimentally, including the dysregulation of specific genes, association of siTPI1 with the downregulation of ‘numerous proliferative pathways’, as well as suggesting a role of *TPI1* in the migration, invasion and apoptosis of the OV56 cell line.

### RNA sequencing analysis supports the predictive accuracy of constraint-based modelling knockdown simulations

As a final validation of constraint-based modelling predictions, the RNAseq data generated from OV56 upon siTPI1 transfection was integrated into an unconstrained copy of the Human1 GEM to model the ‘real-world’ *TPI1* knockdown. In addition, NT siRNA gene expression data was used to constrain an ‘NT siRNA-specific’ model, and this was used as a comparator platform for gene deletion simulations. Given that the aim of the siTPI1 experiment was to reduce the expression of *TPI1* as much as possible, the models to be compared with this ‘real-world’ *TPI1* knockdown model were simplified to full gene deletions. The workflow for this analysis included simulating the deletion of (*n*=147) essential genes in the NT siRNA model, to generate individual *in silico* gene deletion models, and comparing these with the ‘real-world’ *TPI1* knockdown model, constrained using experimental gene expression data from the *in vitro* knockdown (***Figure 6****a*). The null hypothesis here was that the “‘real world’ siTPI1 metabolic model is not more similar to the *in silico TPI1* gene deletion model than the other *in silico* gene deletion models”. A balanced accuracy score from scikit-learn (Pedregosa et al., 2011) was used to calculate the similarity between gene deletion models and the ‘real-world’ *TPI1* knockdown model (*Figure 6a*), and a histogram of these balances accuracy scores was plotted (***Figure 6****b*), so that the null distribution of accuracy scores could be studied.

**Figure 6.**
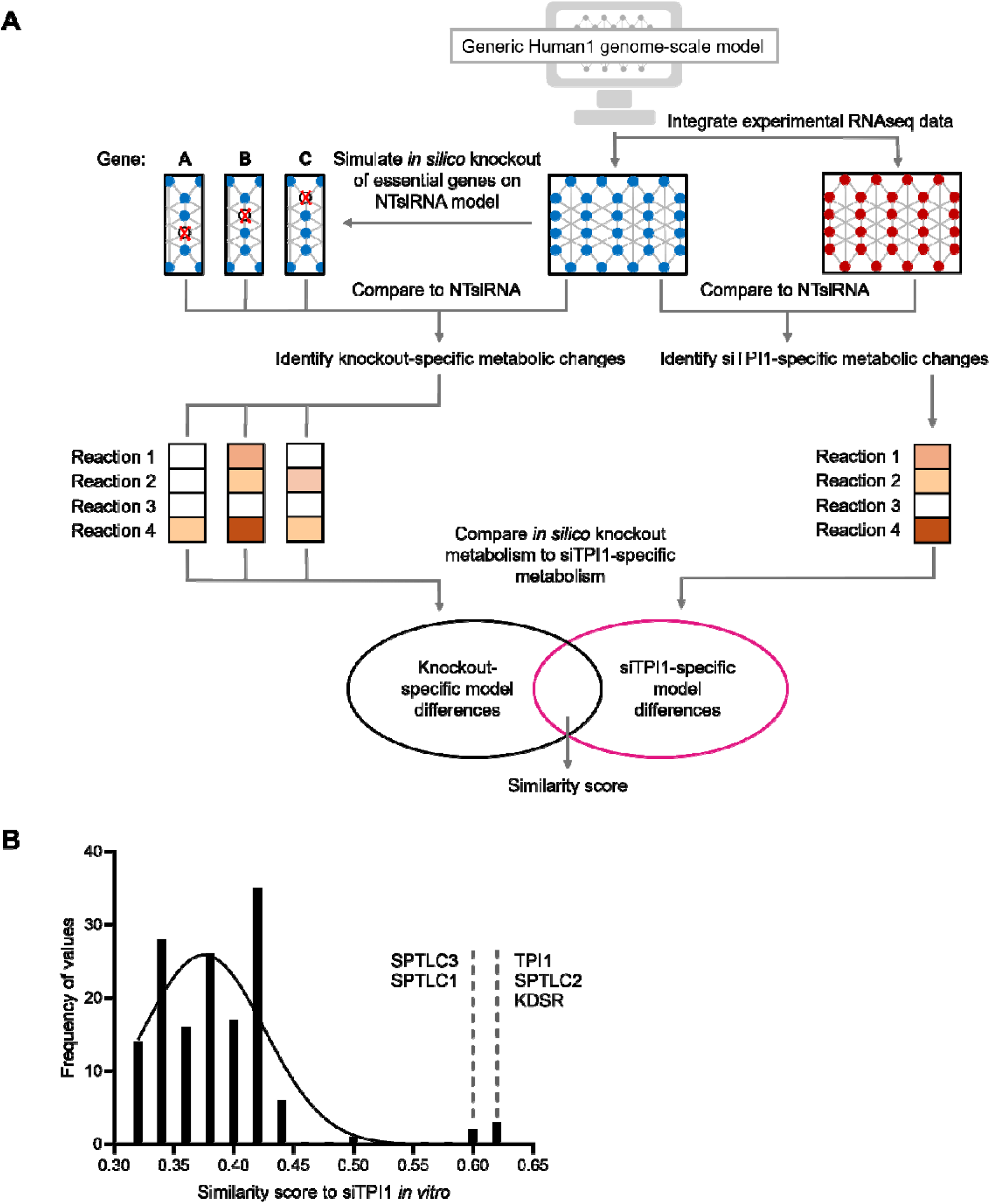
Experimental design and results of knockout simulation validation. A) Experimental design for validation of model knockout simulations. The Human1 GEM was constrained to NTsiRNA and siTPI1-specific models, using experimental RNAseq data. The *in silico* deletion of essential genes was simulated on the NTsiRNA model and compared with the siTPI1-specific model fluxes, using a balanced accuracy score. Gene deletions were performed using COBRApy and essential genes were calculated using MEWpy. B) Histogram to show the similarity between the individual knockout simulation models and the siTPI1-specific model. The five gene knockouts which were the most similar to the siTPI1-specific model are highlighted: TPI1, SPTLC2, KDSR, SPTLC3 and SPTLC1.

Once integrated, gene expression data for the knockdown of *TPI1* in OV56 was able to provide 99.8% coverage of all Human1 genes. When accuracy scores were compared between *in silico* gene deletion models and the ‘real-world’ siTPI1 knockdown model, the highest similarity (accuracy score of 0.619, compared with a mean of 0.375) was obtained between the ‘real-world’ siTPI1 model and the *in silico TPI1* deletion model, meaning the null hypothesis could be rejected (***Figure 6****b*). Other genes that when deleted *in silico* closely resembled the ‘real-world’ siTPI1 model were serine palmitoyltransferase long chain base subunits 1,2 and 3 (*SPTLC1*, *SPTLC2* and *SPTLC3*) and 3-ketodihydrosphingosine reductase (*KDSR*), with scores of 0.604, 0.616, 0.609 and 0.610, respectively (***Figure 6****b*). These genes encode enzymes regulating sphingolipid biosynthesis and metabolism – similar metabolic processes to those that *TPI1* is associated with – since *TPI1* is implicated in glycerophospholipid metabolism via its reactant dihydroxyacetone phosphate (DHAP). These observations further solidify the model predictions, showing the constraint-based models can not only predict the growth effect of a gene knockdown, but also the metabolic flux distribution.

## DISCUSSION

Using a combination of wet-lab experimental and computational approaches, we have built multiple constraint-based models for ovarian cancer cell lines. These models have been shown to accurately reflect the *in vitro* measured growth rates of a panel of ovarian cell lines and *in silico* knockout simulations were supported by a publicly available, CRISPR-Cas9 derived dataset.

Following model simulations, *TPI1* emerged as a top target for inhibiting cell growth. Subsequently, model predictions were validated with experimental long- and short-term proliferation assays. RNAseq analysis identified key signalling pathways and biological functions associated with TPI1, suggesting how this enzyme could regulate the growth of an ovarian cell line. A comparison between computational knockout models and a ‘real-world’ siTPI1 model suggested that constraint-based modelling can accurately predict the metabolic impact of gene knockdowns. Most importantly, this workflow has a translatable, wide-reaching impact, demonstrating how constraint-based models could identify novel targets for the inhibition of cancer cell growth.

### A proposed mechanistic model for the role of TPI1 in ovarian cancer cells

The main catalytic function of TPI1 is the isomerisation of glyceraldehyde 3-phosphate (G3P) and DHAP in glycolysis (Reynolds et al., 1971), which has downstream implications in lipid metabolism and the pentose phosphate pathway (Olivares-Illana et al., 2017). In the context of ovarian cancer, TPI1 is relatively under-researched compared with other key glycolytic enzymes, such as hexokinase 2 (HKII), lactate dehydrogenase (LDH) and pyruvate dehydrogenase kinase 1 (PDK1) (Siu et al., 2019; Xiang et al., 2018; Zhang et al., 2019). One of the few studies exploring TPI1 expression in ovarian cancer found it was more highly expressed in metastatic tumours, specifically brain metastasis, than in the primary ovarian tumour (Yoshida et al., 2013), but no mechanism for this observation has been explored. Therefore, any mechanism that we could propose here would be novel.

Given the limited literature in ovarian cancer, we have evaluated the role of TPI1 across other cancer types (*Figure 7a*) (T. Chen et al., 2017; Jiang et al., 2017; Jin et al., 2022; Li et al., 2023; Ouyang et al., 2018). Among the cancer types studied, TPI1 overexpression has generally been associated with tumour promoting effects, except in hepatocellular carcinoma, where it has been reported to act as a tumour suppressor (Jiang et al., 2017). Several signalling pathways have been associated with the role of TPI1 in regulating cell proliferation, invasion and survival, including the MAPK3/MAPK1 pathway (Li et al., 2023), CDCA5 (T. Chen et al., 2017; Jin et al., 2022), EGFR/MDM2 (Ouyang et al., 2018), the cadherin switch (Jin et al., 2022; Li et al., 2023) and integrin signalling (Q.-Y. Liu et al., 2006).

**Figure 7.**
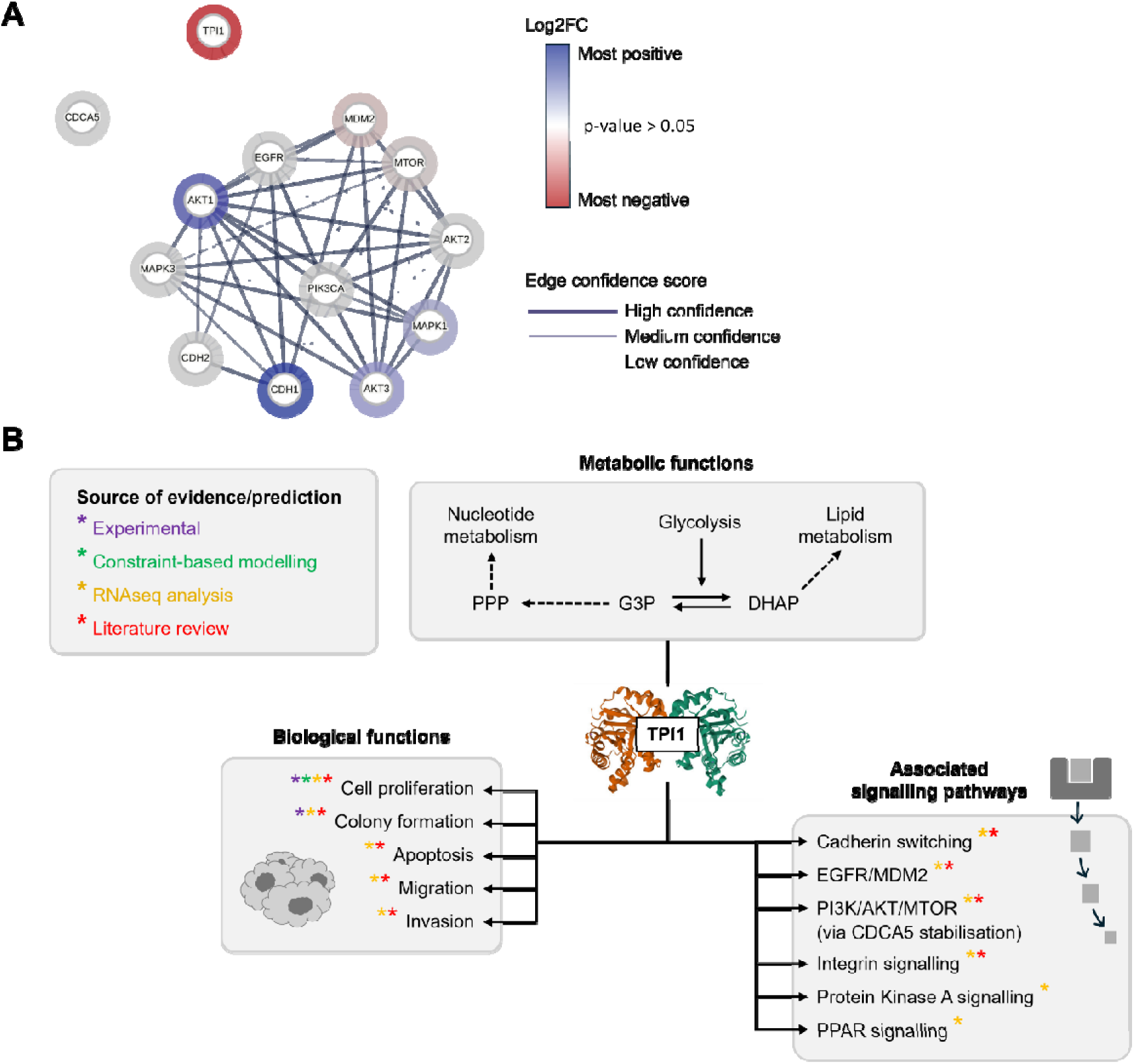
A proposed mechanism for the role of TPI1 in ovarian cancer cells. A) Protein signalling pathways through which TPI1 mediates proliferative processes. This network is speculative, containing information from analysis performed here, and a pan-cancer literature review. Network generated on STRING, using the following parameters: line thickness indicates the strength of supporting data; colour of node indicates differential regulation (between NTsiRNA and siTP1; Log2FC) from RNAseq analysis (white nodes inform no statistical significance (p-value>0.05); active interaction sources contributing to the confidence score were text mining, experiments and databases; low confidence interactions and stronger have been shown. B) An illustration of the metabolic, biological function and downstream pathway associations that work performed here and literature analysis indicate for TPI1, in the context of ovarian cancer. Mechanism is speculative. Literature references are included in-text and do not refer to ovarian cancer specifically, but other cancer types. Dashed arrow indicates metabolite feeding into subsystem. Asterisk indicates data source. Crystal structure of TPI1 downloaded from RCSB Protein Data Bank, accession code 4POC.

Concerning our own results, RNAseq measurements show that *TPI1* knockdown regulates the gene expression of several molecules in signalling pathways that have been associated with TPI1 in other cancer types (*Figure 7a*). Combined with our experimental data indicating roles in cell proliferation and colony formation, we propose a speculative mechanistic model for TPI1 in ovarian cancer (*Figure 7b*). Within this model, we evaluated specific gene-gene interactions to propose signalling pathways that may mediate the activity of TPI1.

Gene expression results highlighted two genes through which *TPI1* could regulate its tumour promoting activity: *CDH1* (encoding E-cadherin protein) and *MDM2* (encoding MDM2 protein). Network analysis identified *CDH1* as central to a gene interaction hub regulating cell proliferation (*Figure 5e*). Within our speculative mechanism, we suggest that the knockdown of *TPI1* leads to increased expression of *CDH1*, based on differential gene expression we observed (*Figure 7a*). Therefore, we propose that reducing the expression of *TPI1* in ovarian cancer would maintain cell-cell adhesion, thereby reducing cell migration and invasion. In addition, the knockdown of *TPI1* led to *MDM2* downregulation (*Figure 7a*), and the MDM2 protein negatively regulates the p53 tumour suppressor. Therefore, in our proposed mechanism, we suggest that if the expression of *TPI1* was reduced, this would reduce the expression of MDM2 and remove negative regulation of p53, allowing p53 to exert its tumour suppressive activity.

For certain signalling pathways, the role of TPI1 was more ambiguous – with gene expression results that could be interpreted as either tumour promoting or tumour suppressing. For example, within the PI3K/AKT/MTOR pathway, there was upregulation of *AKT1* and *AKT3* in response to *TPI1* knockdown, whilst *MTOR* was downregulated (*Figure 7a*). This pattern of regulation is challenging to interpret as individual molecules within the PI3K/AKT/MTOR pathway could have a dual role in cancer signalling, for example, expression of AKT1 or AKT3 is a favourable prognostic marker in renal cancer, but increased expression is an unfavourable marker in liver cancer (AKT1) and stomach cancer (AKT3) (*Human Protein Atlas*, 2024; Uhlén et al., 2015). We found similar counter-intuitive results for the *MAPK1,* which activates the expression of growth-promoting genes in the MAPK/ERK pathway. Work has shown that MAPK/ERK pathway genetic alteration been associated with improved survival in low-grade serous ovarian carcinoma (Manning-Geist et al., 2022). Our results showed increased expression of *MAPK1* upon *TPI1* knockdown (*Figure 7a*), however our assays demonstrated that this correlated with decreased cell proliferation. Overall, these findings point to the adaptive, multi-faceted nature of ovarian cancer gene expression and the potential for the transcriptome to compensate for the knockdown of *TPI1* with the differential regulation of tumour promoting genes. Furthermore, this suggests that future studies should further explore the *TPI1* interactions in ovarian cancer, since our results suggest that its inhibition could be tumour suppressive and underpinned by complex gene networks.

### Translating this workflow to other disease contexts

As has been demonstrated here, the constraint-based modelling of cancer cell lines is a powerful tool for predicting gene targets to inhibit cell proliferation. This approach has proved promising for many other cancer types, for example, facilitating the metabolic subtyping of endometrial cancer (Srivastava & Vinod, 2023), suggesting metabolic features to distinguish intestinal and diffuse-type gastric cancers (Nam & Lee, 2022) and to propose subtype-specific essential genes in colorectal cancer (Schultz & Qutub, 2016). Aside from subtyping, constraint-based modelling has been employed in drug repurposing projects, for example, to predict which existing drugs could potentially reverse gene expression changes associated with prostate cancer (Turanli et al., 2019).

However, the power of constraint-based modelling does not solely apply to cancer research – it has a wide scope across systems biology. To name a couple of applications, constraint-based modelling has been applied to study antibacterial resistance (Chung & Chandrasekaran, 2022; Presta et al., 2017), as well as having been extrapolated to other subject areas, including chemical engineering, where it is a tool to optimise the production of monoclonal antibodies (Hefzi et al., 2016; Park et al., 2024). These examples demonstrate the range of the predictive ability of constraint-based modelling, and combined with the workflow presented here, serve as a framework for future discovery projects.

## Concluding remarks

Here, multiple ovarian metabolic models have been constrained and have served as the framework for hypothesis-generating, discovery biology. Computational gene engineering simulations highlighted TPI1 as a potential target for the inhibition of ovarian cancer cell line growth, and experimental survival and proliferation assays were used to validate these predictions. Notably, gene expression data suggested that knockdown of *TPI1* could downregulate the E-cadherin switch and the MTOR and MDM2 proteins, providing a molecular basis for the dependency of an ovarian cancer cell line on TPI1 for proliferation, thus inspiring future work. Importantly, when RNAseq data originating from the experimental silencing of *TPI1* was fed back into the Human1 GEM as reaction constraints, gene deletion simulations on an NT siRNA control-specific GEM more accurately represented the metabolism of the ‘real-world’ *TPI1* knockdown, than simulations of knockdown of any other enzyme-encoding gene. Given the heterogeneous nature of ovarian cancer, future studies should evaluate differences in regulation and function of TPI1 between the subtypes of ovarian cancer. It will be exciting to see how this workflow is applied by different groups to predict gene targets in cancer metabolism and across systems biology.

## MATERIALS AND METHODS

## BIOINFORMATICS METHODS

### Constraint-based modelling

All coding was performed in Python. Panda data analysis functions and manipulation tools were used (version 1.4.3) (McKinney, 2010; The pandas development team, 2020), as well as SciPy (Virtanen et al., 2020) and scikit-learn (Pedregosa et al., 2011). The single-omics integration algorithm used for constraint-based modelling is available on the accompanying Github repository (https://github.com/katemeeson/PhD_2024) (Meeson & Schwartz, 2024). This algorithm was applied to the Human1 generic GEM (Robinson et al., 2020) (https://github.com/SysBioChalmers/Human-GEM; 2021) and uses FBA, with biomass production as the objective function. The COBRA and MEWpy toolboxes were used for constraint-based simulations, including gene deletions and FBA (Ebrahim et al., 2013; Pereira et al., 2021). The Gurobi solver 11.0 was used for optimisation (Gurobi Optimization, LLC, 2023).

### Input datasets

Datasets for the constraint of Human1 GEM and its validation were accessed from the Cancer Cell Line Encyclopedia (CCLE) and have been described in detail in Supplementary Table 1. Transcriptomics measurements were used to constrain the Human1 GEM and a CRISPR-Cas9 derived gene dependency dataset (DepMap) was used to validate gene deletion predictions, both of which were accessed via a database portal (https://depmap.org/portal/) (Barretina et al., 2012; Ghandi et al., 2019). Cell line annotations, including the source of experimental growth rates and the optimal media conditions, which were replicated *in silico*, have been described in Supplementary file, entitled ‘CCLE_2020_transcriptomics_annotations.xlsx’.

### RNA sequencing

Total RNA was isolated using the RNeasy mini kit (QIAgen; Cat #74104#). RNA sequencing was performed by Genomic Technologies Core Facility at the University of Manchester. Quality and integrity were assessed using a 4200 TapeStation (Agilent Technologies) and libraries generated using the Illumina® Stranded mRNA Prep (Illumina, Inc). Polyadenylated mRNA was purified from total RNA isolates, using poly-T, oligo-attached magnetic beads. The mRNA was fragmented under elevated temperature and reverse transcribed into first strand cDNA using random hexamer primers and Actinomycin D. Second strand cDNA was synthesized to yield blunt-ended, double-stranded cDNA fragments. Strand specificity was maintained by dUTP-incorporation. Following a single adenine base addition, adapters with a corresponding, complementary thymine overhand were ligated to the cDNA fragments to prepare for dual indexing. PCR amplification was used to add the index adapter sequences to create the final cDNA library. Libraries were multiplexed and pooled prior to loading onto the appropriate flow-cell. The flow-cell was paired-end sequenced (59+59 cycles, plus indices) on an Illumina NovaSeq6000 instrument. Output was demultiplexed and BCL-to-FastQ conversion was performed using Illumina’s bcl2fastq software (2.20.0.422). Quality of stranded paired-end RNAseq reads was assessed using FastQC (v.0.11.3) and FastQ Screen (v0.14.0) (*Babraham Bioinformatics - FastQ Screen*, 2024). BBDuk was used for adapter and low-quality base trimming (BBMap suite v38.96) (*BBMap*, 2023). Trimmed reads were mapped against HG38 and genes annotation was performed using Gencode (v42) using STAR (‘-quantModeGeneCounts’ option) (v2.7.10a) (Dobin et al., 2013). DESeq2 analysis (v1.26.0) was used to identify differentially expressed genes, with alpha=0.05 (Love et al., 2014), with the lfcShrink function (with apeglm method) applied. Media-of-ratios method was used for normalization of raw counts (Anders & Huber, 2010). Rlog function was used to obtain regularized logarithm expression values. The R package pheatmap was used to produce heatmaps (v1.0.12) (Kolde, 2019) and PCA was performed using the prcomp function from the built-in R package stats.

### Functional enrichment of RNAseq DESeq2 results

QIAGEN Ingenuity Pathway Analysis (IPA) was used for functional enrichment of differentially expressed genes (QIAGEN Inc., https://digitalinsights.qiagen.com/IPA) (Krämer et al., 2014). For all IPA analysis parameters were as follows: direct and indirect relationships were included; interactions and causal network selected; all data sources included; experimentally observed or high (predicted) miRNA confidence were filtered; Human species only; all node types except for ‘fusion gene/product’; all tissues, cell lines and mutations. Statistical parameters for functional enrichment were as follows: Log2FC less than or equal to -0.5 or greater than or equal to 0.5; false discovery rate (FDR) of less than 0.01. Additional settings for functional enrichment ‘Disease and Functions’ analysis was a z-score of ≥ +2 or ≤ -2 (activation or inhibition of a pathway, respectively). These z-score thresholds have been advised by QIAGEN to indicate significance (QIAGEN, 2024). The date of IPA analysis was 05/12/2023. STRING protein-protein database was used to infer a speculative mechanistic model for the gene network interaction surrounding TPI1 in OV56 (Szklarczyk et al., 2023); parameters included in figure legend.

## IN VITRO METHODS

### Cell culture

The A549 human lung carcinoma cell line (ATCC, Cat#CCL-185; RRID: CVCL_0023) and OV56 serous adenocarcinoma (Sigma Aldrich, Cat# 96020759; RRID: CVCL_2673) cell lines were cultured in their optimal media conditions, in a humidified environment with 5% CO_2_, at 37°C. Experimental conditions for cell culture have been described in Supplementary Table 3.

### siRNA transfection

OV56 and A549 cells were transfected with ON-TARGETplus SMARTpool Human TPI1 and MYC siRNA (Horizon Discovery). A final concentration of 0.66 µM of siRNA was added to cells for transfection in a 24-well plate (to study cell proliferation), and cells were incubated for 48 hours at a seeding density of 40,000 cells/well. To generate samples for RNA extraction, a seeding density of 1,300 cells/well was incubated with siRNA for 48 hours and a final concentration of 66 nM siRNA was achieved. Details of siRNA sequences have been described in Supplementary Table 4.

### Colony formation assays

A549 and OV56 cells were seeded at 250–350 cells/well in a 6-well plate. Cells were incubated for 10 days with siRNA added to cells immediately upon plating and media was refreshed at the halfway point of the experiment. Once colonies had formed, cells were washed in PBS and fixed in 1% Formaldehyde and stained with 0.05% (w/v) Crystal Violet solution. Quantification of colony formation samples was performed on the VARIOSKAN LUX Colorimeter (Thermo Fisher Scientific) at a wavelength of 590 nm.

### Incucyte live cell imaging and growth ratio calculations

Label-free phase images were taken of cells every two hours across three fields of view per well using the cell-by-cell Incucyte S3^®^ software on the Incucyte S3^®^ live-cell analysis system (Sartorius), with a 20X objective. Therefore, over a 120-hour period, there were 60 values calculated. Prism 10 (GraphPad) was used for visualization of quantification and statistical analysis. For the calculation of the ‘growth ratio’ which was used to compare the growth of cells before and after siTPI1 transfection, the doubling time before transfection was divided by the doubling time after siRNA transfection. In this way, a ‘growth ratio’ of less than 1 referred to a growth inhibition effect and a ratio greater than 1 indicated siRNA transfection increased cell proliferation.

### Immunoblotting

Following culture, cells were washed in PBS and harvested using trypsin and cell scraping. Protein was extracted using 6X SDS buffer and boiled for 5 minutes at 100°C. Cells were loaded onto and separated using a NuPAGETM 4-12% (v/v) Bis-Tris protein gel (1.0mm; Life Technologies). Proteins resolved using SDS-PAGE were transferred to a methanol-activated Immobilon-P membrane, (Merck Millipore) through electroblotting at 50V for 60 minutes, using the Mini-PROTEAN® Tetra System (BIO-RAD) in 1X transfer buffer (25mM Tris, 190mM glycine, 0.1% (w/v) SDS, 20% (v/v) methanol). Membranes were blocked with 5% (w/v) dried skimmed milk (Marvel) made up with TBS-T and incubated overnight at 4°c with primary antibodies: rabbit polyclonal anti-TPI1 (Proteintech, sc-166785, 1:1000 dilution) and sheep polyclonal anti-TAO1 (1:1000 dilution) (Westhorpe et al., 2010). After incubation, membranes were washed three time in TBS-T then incubated in horseradish-peroxide (HRP)-conjugated secondary antibodies for 2 hours. Membranes were imaged using EZ-Chemiluminescence (Geneflow ltd) and a ChemiDocTMTouch Imaging System (Bio-Rad) and Adobe Photoshop® CC 2024 (Adobe Systems Inc.) were used for image processing.

### Quantification and statistical analysis

Prism 10 (GraphPad) was used for statistical analysis, where *p<0.05, **p<0.01, ***p<0.001, ****p<0.0001, ns: p>0.05. Details of statistical analyses are described in the figure legends.

## DATA AVAILABILITY

Gene expression data presented here will be deposited with EMBL-EBI upon publication.

## AUTHOR CONTRIBUTION STATEMENT

All authors contributed to the study conception and design. Material preparation, data collection and analysis were performed by Kate Meeson. The first draft of the manuscript was written by Kate Meeson and all authors commented on versions of the manuscript. All authors read and approved the final manuscript.

○ Methodology: Meeson, Schwartz and Taylor
○ Investigation: Meeson
○ Data curation: Meeson
○ Validation: Meeson, Schwartz and Taylor
○ Formal Analysis: Meeson
○ Conceptualisation: Meeson, Schwartz and Taylor
○ Funding acquisition: Schwartz and Taylor
○ Supervision: Schwartz and Taylor
○ Project administration: Meeson, McGrail, Schwartz and Taylor
○ Resources: Taylor
○ Software: Meeson and Schwartz
○ Visualisation: Meeson
○ Writing original draft: Meeson
○ Writing (review and editing): Meeson, McGrail, Schwartz and Taylor

## DECLARATIONS

The authors have no competing interests to declare that are relevant to the content of this article.

## ACKNOWLEDGEMENTS

For the duration of this study, J-M.S and K.M were funded by the Medical Research Council (MRC) awarded to J-M.S. K.M was funded by the MRC-DTP training programme (MR/N013751/1). S.S.T is funded by a Cancer Research UK Programme [C1422/A31334 to S.S.T.]; Medical Research Council [MR/X008088/1 to S.S.T.]; NIHR Manchester Biomedical Research Centre [NIHR203308]; Cancer Research UK Manchester Centre [C147/A25254]; the views expressed are those of the author(s) and not necessarily those of the NIHR or the Department of Health and Social Care. We thank members of the Taylor lab for experimental advice and comments on the manuscript, specifically Anthony Tighe, Camilla Coulson-Gilmer and Samantha Littler. We thank the Genomic Technologies Core Facility at the University of Manchester for running the RNAseq and initial analysis. Data used for model constraint has been made publicly available, thanks to the Cancer Cell Line Encyclopedia.

## Notes

https://github.com/katemeeson/PhD_2024

### Competing Interest Statement

The authors have declared no competing interest.

