## Supplemental Tables and Figure for "Transcriptome-Driven Constraint-Based Modelling Reveals Metabolic Targets for Ovarian Cancer"

**Supplementary data**

| **Dataset** | **General description** | **Normalisation and processing details** | **Reference and accession** |
| --- | --- | --- | --- |
| **CCLE transcriptomics** | 64 ovarian cell lines;  53,970 genes | RSEM; Log_2_(TPM+1) | CCLE_expression.csv; Depmap |
| **CCLE CRISPR-Cas9 gene dependency dataset** | 1,086 total cell lines; 17,387 genes |  | CRISPR_gene_dependency.csv; Depmap |

**Table 1. Description of input and validation datasets.** RSEM: RNAseq by expectation-maximisation; TPM: transcripts per million. Cell line annotations including source of experimental growth rate and optimal media conditions, which were replicated in silico, have been described in the Supplementary file ‘CCLE_2020_transcriptomics_annotations.xlsx’.

| **Cell line model** | **p-value** | **r** |
| --- | --- | --- |
| **COV504** | 7.6398 x 10^-22^ | -0.498762 |
| **SNU8** | 6.6205 x 10^-52^ | -0.468450 |
| **JHOS2** | 3.00298 x 10^-31^ | -0.452691 |
| **PEA1** | 1.63158 x 10^-97^ | -0.404164 |
| **59M** | 6.41651 x 10^-10^ | -0.368615 |
| **OAW28** | 9.09970 x 10^-79^ | -0.355036 |
| **PEO4** | 5.18932 x 10^-9^ | -0.183914 |
| **CAOV3** | 2.68982 x 10^-5^ | -0.173970 |
| **JHOS4** | 4.09742 x 10^-4^ | -0.165705 |
| **ES2** | 8.57693 x 10^-10^ | -0.139105 |
| **PEO1** | 2.68663 x 10^-9^ | -0.130448 |
| **CAOV4** | 2.94089 x 10^-10^ | -0.119685 |
| **NIHOVCAR3** | 0.0452459 | -0.117682 |
| **COV318** | 0.1400247 | -0.100973 |
| **HEYA8** | 0.6149733 | -0.028539 |
| **KURAMOCHI** | 0.6070811 | 0.012544 |
| **COV362** | 0.3106583 | 0.023102 |

**Table 2. Pearson correlation and p-values for gene knockdown simulations.** The 13 out of 17 significant models are those below with p≤0.05. Ordered from smallest to largest r.

| **Cell line** | **Experimental growth conditions** | **Seeding dilution into new flask** |
| --- | --- | --- |
| **A549** | RPMI1640 + 10% fetal bovine serum (FBS) + 50 µg/mL penicillin/streptomycin | 1:6 |
| **OV56** | 1:1 ratio of Dulbecco’s Modified Eagle’s Medium (DMEM) and Ham’s F-12 nutrient mix + 5% FBS + 2 mM glutamine + 0.5 μg/mL hydrocortisone (Sigma Aldrich) + 10 μg/mL insulin (Sigma Aldrich) + 50 μg/mL penicillin/streptomycin | 1:8 |

**Table 3. Experimental conditions for cell culture.** All media and supplements are from Life Technologies.

| Target gene | Manufacturer details | Sequences |
| --- | --- | --- |
| TPI1 | ON-TARGETplus Human TPI1 siRNA  Cat # L-009776-00-0005 | 1. GAGCCUGUGUGGGCCAUUG 2. CCAGGAAGUACACGAGAAG 3. GGGUGGUGCUUCCCUCAAG 4. GCAGAAAGUGGCCCAUGCU |
| MYC | ON-TARGETplus MYC siRNA oligonucleotide #4  Cat# J-003282-26  (Topham et al., 2015) | CGAUGUUGUUUCUGUGGAA |
| Non-targeting control pool | ON-TARGETplus Non-targeting Control Pool  Cat# D-001810-10 | 1. UGGUUUACAUGUCGACUAA 2. UGGUUUACAUGUUGUGUGA 3. UGGUUUACAUGUUUUCUGA 4. UGGUUUACAUGUUUUCCUA |

**Table 4. Details of siRNA sequences.**

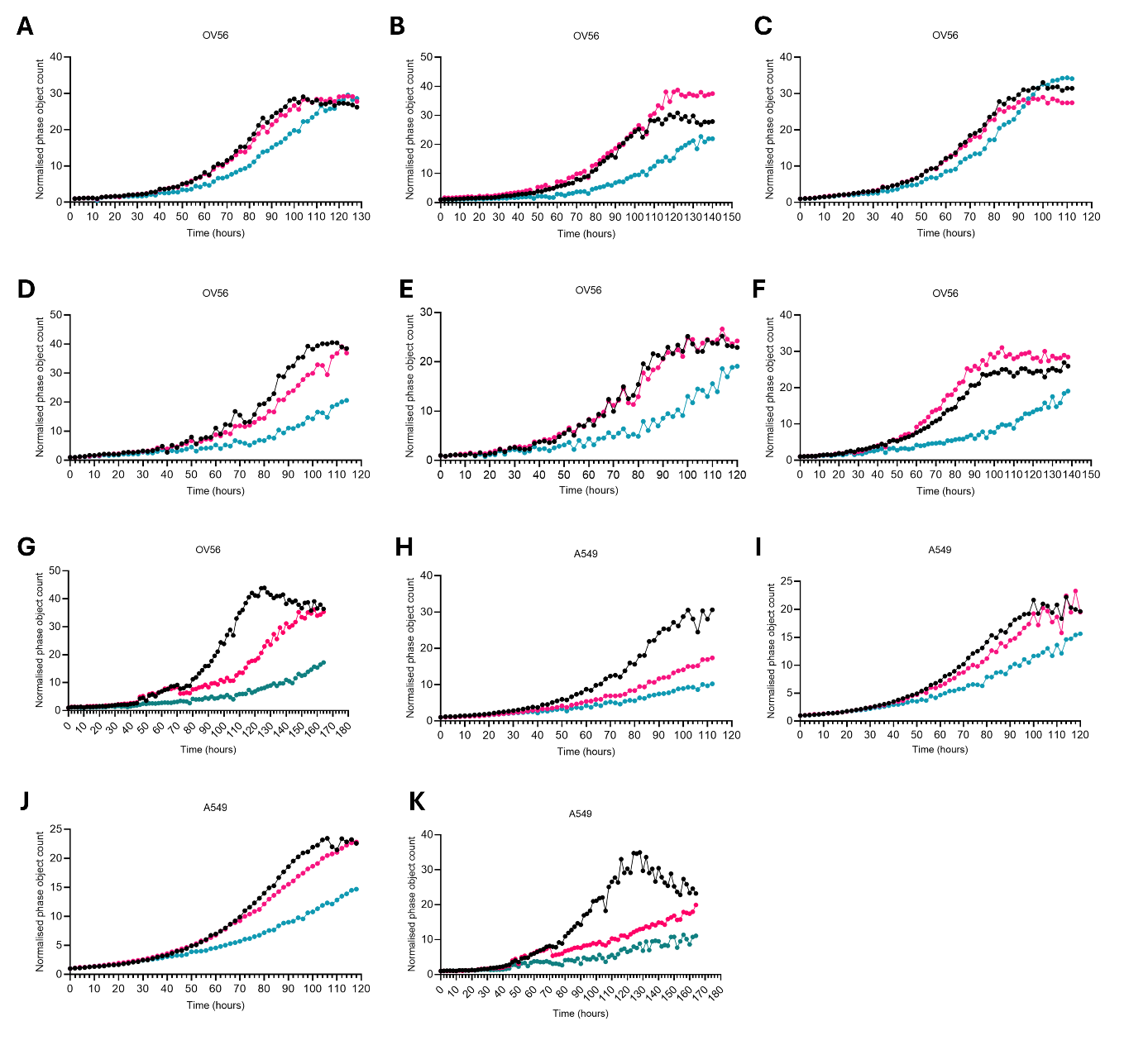

**Figure S1. Biological repeats for the siRNA transfection of OV56 and A549.**

**A-G** Growth of OV56 cells upon siRNA transfection (NTsiRNA: black, siTPI1: pink, siMYC: teal). Live cell microscopy assay, using the Incucyte S3^®^ live-cell analysis system (Sartorius). **H-K** Growth of A549 cells upon siRNA transfection (NTsiRNA: black, siTPI1: pink, siMYC: teal). Live cell microscopy assay, using the Incucyte S3^®^ live-cell analysis system (Sartorius). Associated with Figure 3, showing biological replicates for the siRNA transfection of OV56 and A549 cell lines. The growth effect, as calculated using the normalised phase object count at 100 hours for siTPI1 transfection compared to NTsiRNA transfection has been included in Figure 3F.
